# c-di-GMP-linked phenotypes are modulated by the interaction between a diguanylate cyclase and a polar hub protein

**DOI:** 10.1101/154005

**Authors:** Gianlucca G. Nicastro, Gilberto H. Kaihami, André A. Pulschen, Jacobo Hernandez-Montelongo, Ana Laura Boechat, Thays de O. Pereira, Eliezer Stefanello, Pio Colepicolo, Christophe Bordi, Regina L. Baldini

## Abstract

c-di-GMP is a major player in the decision between biofilm and motile lifestyles. Several bacteria present a large number of c-di-GMP metabolizing proteins, thus a fine-tuning of this nucleotide levels may occur. It is hypothesized that some c-di-GMP metabolizing proteins would provide the global c-di-GMP levels inside the cell whereas others would maintain a localized pool, with the resulting c-di-GMP acting at the vicinity of its production. Although attractive, this hypothesis has yet to be proven in *Pseudomonas aeruginosa.* We found that the diguanylate cyclase DgcP interacts with the cytosolic region of FimV, a peptidoglycan-binding protein involved in type IV pilus assembly. Moreover, DgcP is located at the cell poles in wild type cells, but scattered in the cytoplasm of cells lacking FimV. Overexpression of DgcP leads to the classical phenotypes of high c-di-GMP levels (increased biofilm and impaired motilities) in the wild-type strain, but not in a *AfimV* background. Therefore, our findings strongly suggest that DgcP is regulated by FimV and may provide the local c-di-GMP pool that can be sensed by other proteins at the cell pole, bringing to light a specialized function for a specific diguanylate cyclase.

## Introduction

Over the past decades, (3'-5')-cyclic diguanylic acid (c-di-GMP) has been characterized as an important second messenger in bacteria. The concentration of c-di-GMP within the cell is associated with cellular behavior: high c-di-GMP levels are linked to biofilm formation and low levels to the motile planktonic lifestyle (Simm *et al.,* 2004; Romling *et al.,* 2013). This molecule is synthesized from GTP by a class of enzymes known as diguanylate cyclases (DGC) bearing a conserved GGDEF motif (Chan *et al.,* 2004). The c-di-GMP hydrolysis reaction is performed by phosphodiesterases (PDE) with EAL or HD-GYP domains, which cleave c-di-GMP to pGpG or GMP, respectively (Schmidt *et al.,* 2005; Ryan *et al.,* 2006). Multiple genes coding for the c-di-GMP-metabolizing proteins are found in a variety of bacterial genomes. A puzzling question in the study of c-di-GMP signaling is how the bacterial cell integrates the contributions of multiple c-di-GMP-metabolizing enzymes to mediate its cognate functional outcomes. Merritt and collaborators showed that the *P. aeruginosa* phenotypes controlled by two different DGC have discrete outputs despite the same level of total intracellular c-di-GMP (Merritt *et al.,* 2010). These data support the model in which localized c-di-GMP signaling contributes to the action of proteins involved in the synthesis, degradation, and/or binding to a downstream target (Romling *et al.,* 2013). Studies of c-di-GMP signaling regulation during the swarmer to stalked-cell transition in *Caulobacter crescentus* also supports this hypothesis. In this dimorphic bacterium, PleD is a DGC that is inactive in swarmer cells and is activated during the swarmer-to-stalked cell differentiation (Aldridge *et al.,* 2003; Paul *et al.,* 2004). Activation of PleD is coupled to its subcellular localization at the stalk pole, suggesting that PleD activates nearby downstream effectors involved in pole remodeling (Paul *et al.,* 2007). Opposite to PleD, the EAL domain protein TipF localizes at the swarmer pole, where it contributes to the proper placement of the motor organelle in the polarized predivisional cell (Davis *et al.,* 2013).

Even though a large body of research on c-di-GMP regulation in *P. aeruginosa* is available, it is still unclear whether compartmentalization of c-di-GMP signaling components is required to mediate an appropriate c-di-GMP signal transduction. The genome of *P. aeruginosa* strain PA14 presents forty genes coding for proteins associated with c-di-GMP metabolism (Kulasakara *et al.,* 2006; Lee *et al.,* 2006). Some of these proteins were already characterized and a few of them present a specific localization within the cell. For instance, the DGC WspR is associated to contact-dependent response to solid surfaces. Activation of the Wsp system by contact leads to the formation of subcellular clusters of WspR followed by synthesis of c-di-GMP, increasing exopolysaccharide production and biofilm formation (Huangyutitham *et al.,* 2013). The DGC SadC is a central player in Gac/Rsm-mediated biofilm formation (Moscoso *et al.,* 2014) and influences biofilm formation and swarming motility via modulation of exopolysaccharide production and flagellar function (Merritt *et al.,* 2007). Recently, it was demonstrated that SadC activity is promoted by membrane association, with the formation of active DGC oligomers (Zhu *et al.,* 2016). The PDE DipA/Pch is essential for biofilm dispersion (Roy *et al.,* 2012) and promotes c-di-GMP heterogeneity in *P. aeruginosa* population (Kulasekara *et al.,* 2013). This PDE is partitioned after cell division and is localized to the flagellated cell pole by the chemotaxis machinery. This asymmetric distribution during cell division results in a bimodal distribution of c-di-GMP (Kulasekara *et al.,* 2013).

Previously, we demonstrated that PA14_72420 is an enzymatically active DGC that increased imipenem fitness (Nicastro *et al.,* 2014). Although this protein has been the subject of recent publications that named it as DgcP (Aragon *et al.,* 2015), its molecular function has not yet been addressed. Thus, we decided to pursue its role by seeking for DgcP interaction partners that could participate in the same signaling pathway. DgcP was found to interact with the inner membrane protein FimV, which has a regulatory role in type IV pilus (T4P) function. Moreover, we determined that DgcP localizes to cell poles in a FimV-dependent manner and is more active when the FimV protein is present. We suggest that the DgcP regulation by FimV may provide a local c-di-GMP pool at the cell pole, making this second messenger available for the c-di-GMP binding proteins that may regulate the machineries associated with the cell motility, such as the flagellum and pili.

## Results

### DgcP interacts with FimV

One of the interesting paradoxes of signal transduction by c-di-GMP is the redundancy of DGCs and PDEs in bacterial cells. It has been shown that distinct phenotypes are controlled by specific DGCs or PDEs (Merritt *et al.,* 2010; Romling *et al.,* 2013). The specificity of DGCs and PDEs has been proposed to be related to protein localization, allowing the regulation of subcellular pools of c-di-GMP close to a target receptor (Merritt *et al.,* 2010; McDougald *et al.,* 2012). Therefore, we sought, using bacterial two-hybrid system, for interaction partners of DgcP that could give a hint of DcgP localization and function. The bait plasmid *pKT25_dgcP* was constructed with the complete *dgcP* coding region (Fig. 1 A) and co-transformed in BTH101 *E. coli* cells with a *P. aeruginosa* fragment prey library cloned in the pUT18C plasmid (Houot *et al.,* 2012). About 100,000 co¬transformants were obtained and 40 positive red colonies identified. All positives clones were retested by individually transforming different pUT18 derivative preys and the pKT25_dgcP or pKT25 empty vector plasmids into BTH101 cells. From the 40 clones initially obtained, 26 were confirmed and the inserts were further identified by DNA sequencing. Only two clones presented *in frame* inserts, one of them corresponding to a short peptide fragment (36 amino acids) of the cytoplasmic, C-terminal portion of FimV (Fig. 1 B, red box). FimV is a large protein (924 amino acids, 97 kDa) containing a periplasmic domain with a peptidoglycan-binding LysM motif connected via a single transmembrane segment to a highly acidic cytoplasmic domain with three predicted protein-protein interaction tetratricopeptide repeat (TPR). FimV is part of the T4P secretion machinery in *Pseudomonas* (Wehbi *et al.,* 2011) and is required for localization of some T4P assembly components to the cell pole (Carter *et al.,* 2017). As T4P are important for initial cell attachment and the *ΔdgcP* mutant is impaired in this trait (Aragon *et al.,* 2015), we decided to further investigate the interaction of FimV with DgcP. Two constructs, one containing the FimV cytoplasmic region and the other, a smaller cytoplasmic fragment were cloned into pUT18 plasmid (Fig. 1 B). After co-transformation of each of these plasmids with pKT25_dgcP in BTH101 cells, we confirmed that DgcP interacts with the cytoplasmic region of FimV (Fig. 1 C and D). Stronger interactions were observed with the constructs containing the prey fragment or the full cytoplasmic region.

**Figure 1.**
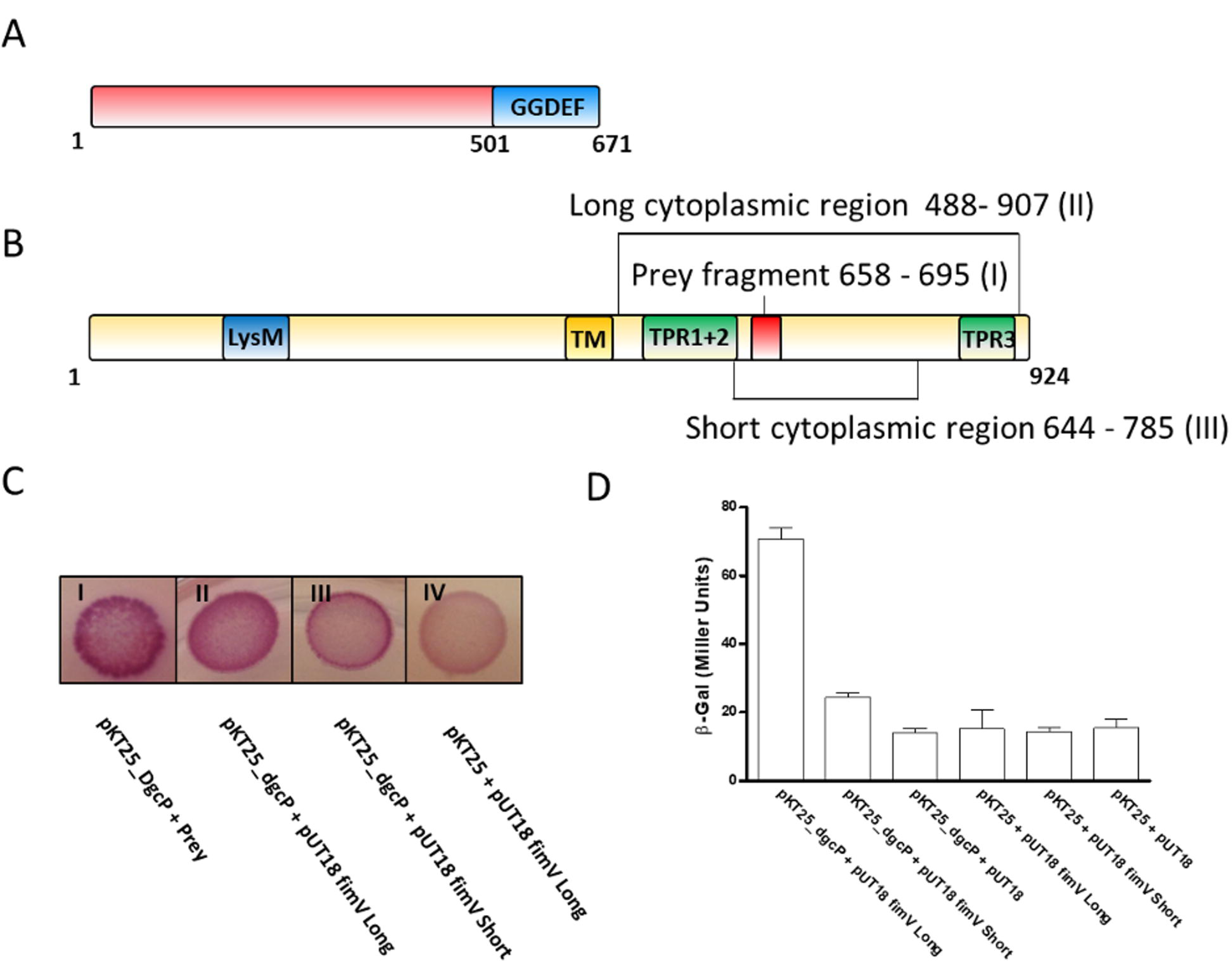
DgcP interacts with FimV. Schematic diagrams of DgcP (**A**) and FimV (**B**), showing the GGDEF domain of DgcP; the transmembrane region (TM), TPR motifs and the LysM domain of FimV. The red box in FimV corresponds to the prey fragment and lines show the regions cloned to confirm the interaction. The *E. coli* host strain BTH101 was cotransformed with pKTA25_DgcP (full length) and pUT18_FimV constructs, as indicated in the figure; the interactions were observed in MacConkey plates as red colonies (**C**) or measured using the (B-galactosidase activity as a reporter. Data are the means ± SD from three replicates (**D**).

### FimV localizes DgcP at the cell poles

Type IV pili are surface appendages that localize at the cell poles in *P. aeruginosa.* There are several polar localized proteins involved in the assembly and regulation of the T4P system and FimV is one of them (Wehbi *et al.,* 2011; Inclan *et al.,* 2016; Buensuceso *et al.,* 2016). Due to its interaction with FimV, we asked whether DgcP would also localize at the cell poles. Indeed, when cells overexpresses the fusion protein DgcP-msfGFP, polar fluorescent foci were observed in virtually 100% of both PA14 (not shown) and *ΔdgcP* strains (Fig. 2 A). This localization is lost when DgcP-msfGFP is expressed in a *ΔfimV* mutant, where the fluorescence is scattered throughout the cells (Fig. 2 B and C). To narrow the DgcP region important to the interaction, we use two different DgcP-msfGFP fusions, one containing the first 199 amino acids (N-ter-DgcP-msfGFP) and the other the last 461 amino acids (C-ter-DgcP-msfGFP) (Fig. 2 A). The C-terminal fusion fluorescence was scattered in the cells (not show) and cells overexpressing the N-ter-DgcP-msfGFP presented a growth defect, precluding its analysis. Therefore, we cannot determine precisely which region of DgcP is important to polar localization.

**Figure 2.**
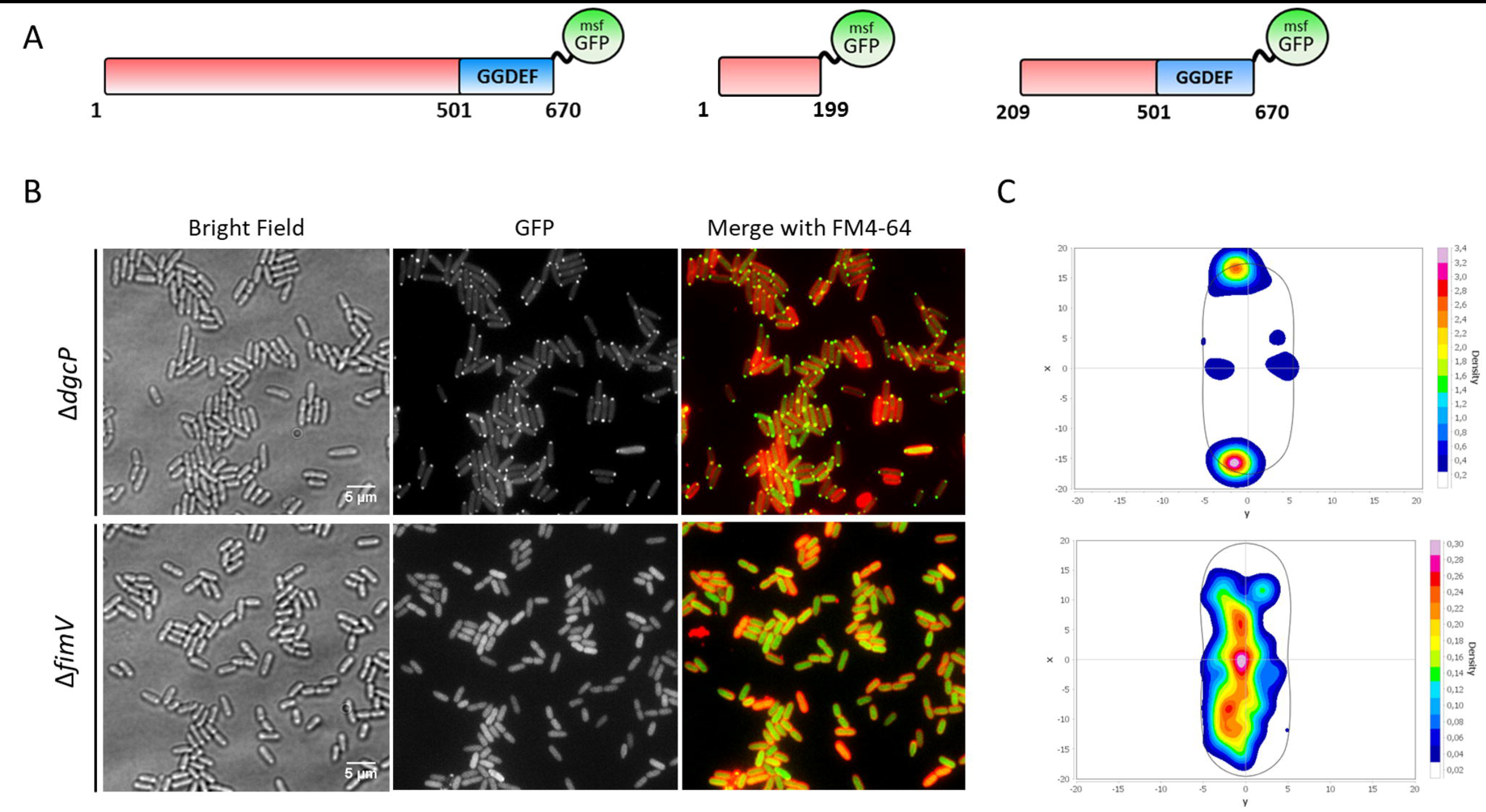
DgcP localizes at the cell poles in a FimV-dependent manner. msfGFP was fusioned to DgcP full-length (1-670), its N-terminal (1-199) or the C-terminal (209-670) regions, as depicted (**A**). The full length protein localizes at the cell poles in the *ΔdgcP* background (B, top panels), but deletion of *fimV* leads to a loss of localization (**B**, bottom panels). The intensity of the GFP fluorescence was measured in 300 cells of each strains and a heat map of DgcP_msfGFP localization was obtained with MicrobeJ, as described in Material and Methods (**C**).

As the DgcP C-terminal construct is not localized and there are no predicted domains in the N-terminal region, we aligned 85 sequences of putative DgcP orthologues from 75 species, including 53 Pseudomonadaceae and 32 other proteobacteria, such as Vibrionales and Alteromonadales (Table S2). We found that the N-terminal region is composed by two separate conserved segments (Fig. S1) and both or one of them may be responsible for interaction with FimV, which homologues are present in all those bacterial genomes.

### DgcP plays a role in biofilm formation

The role of DgcP in biofilm formation has been investigated by different groups under different conditions. Kulasekara and collaborators showed that mutation in DgcP abolished biofilm formation in LB medium (Kulasakara *et al.,* 2006) and Ha and collaborators showed that a *dgcP* mutation did not affect biofilm formation in M63 minimum medium (Dae-Gon Ha *et al.,* 2014). Aragon and collaborators demonstrated that deletion of *dgcP* orthologues in *Pseudomonas savastanoi* pv. savastanoi and *P. aeruginosa* PAK indeed decreased biofilm formation in LB (Aragon *et al.,* 2015). Here, we confirmed that the PA14 *ΔdgcP* mutant is impaired in biofilm formation in the rich medium LB, but minor differences were observed in minimal M63 medium (Fig. 3 A). These results are in agreement with the differences observed by the two previous studies (Kulasakara *et al.,* 2006; Dae-Gon Ha *et al.,* 2014). The expression *in trans* of *dgcP* restores the phenotype of biofilm defect on *ΔdgcP* (Fig. 3 B and C). In LB, *ΔdgcP* was not able to form a biofilm and just a few adherent cells were observed by confocal laser scanning microscopy (CLSM) after 16h post¬inoculation, while the wild type PA14 biofilm was more developed at this time point (Fig. 3 C). At 72 hours, *ΔdgcP* biofilm was thin and undifferentiated (Fig. 3 D). As FimV is also involved in biofilm formation (Fig. 5), this is another indication that they have complementary roles in the cells, probably related to the T4P function.

**Figure 3.**
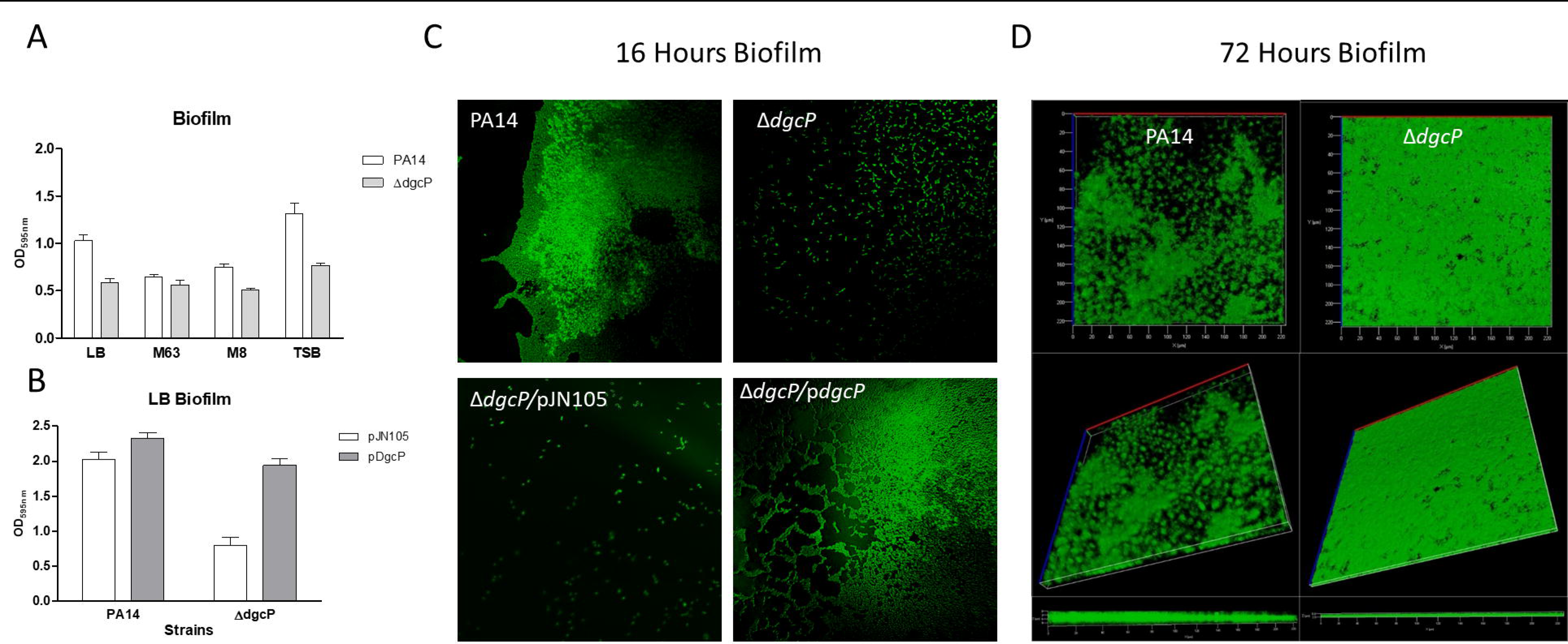
Mutation in ***dgcP*** affects biofilm formation. PA14 and the *ΔdgcP* strains were inoculated at OD_600_□=□0.05 in 48 well polystyrene plates with the media shown and kept at 30°C for 16Lh without agitation. The medium was discarded and adherent cells were washed and stained with 1% crystal violet, washed and measured at OD_595_. (A). The same procedure was carried out for the strains overexpressing DgcP in LB with 0.2% arabinose (B). 3D pictures resulting from CLSM after 16 h (C) and 72 h (D) of biofilm formation in LB at 30^o^C in an 8-well Lab-tek chambered coverglass system. Data in A and B are the means ± SD from five replicates.

**Figure 5.**
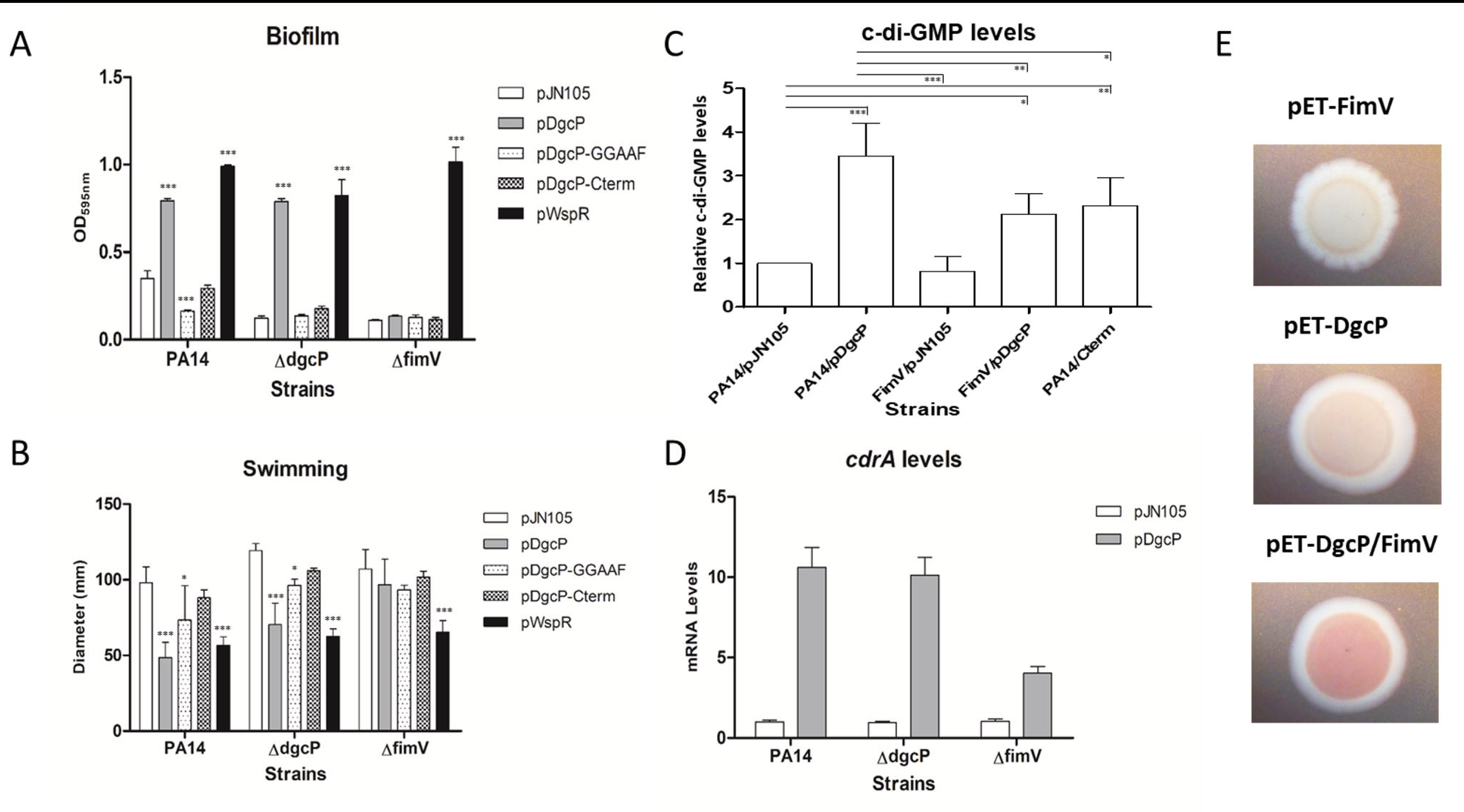
DgcP activity is FimV dependent. Full-length wild type DgcP (pDgcP) or DgcP mutated in its GGDEF domain (pDgcP-GGAAF), only DgcP C-terminal wild-type region (pDgcP-Cterm) and WspR (pWspR) were overexpressed from the pJN105 vector in PA14, *ΔdgcP* and *ΔfimV* backgrounds. Biofilm (A), swimming motility (B), c-di-GMP quantitation (C) and *cdrA* mRNA relative levels (D) were assayed. FimV, DgcP or both were expressed in *E. coli* and the EPS production was assessed in Congo red plates (E). Data are the means ± SD from three replicates. *, p<0.05; **, p<0.01; ***, p<0.001

### DgcP has a role in twitching motility

*P. aeruginosa* utilizes T4P to move across solid surfaces in a process known as twitching motility. As T4P are regulated by FimV, we decided to investigate if DgcP is important for twitching. A small portion of the outer edge of the bacterial streak was taken and stabbed into the bottom of the agar plate or placed on a thin layer of solidified media and covered with a glass coverslip. Cells were incubated and active colony expansion occurred at the interstitial interface. Twitching motility was analyzed by staining the plates with crystal violet after 16h (Fig. 4 A and B) or by phase contrast microscopy after 4 hours of colony expansion (Fig. 4 C). As expected, the *ΔdgcP* mutant presented decreased twitching motility with a less defined structure whereas PA14 presented a well-defined lattice-like network. The *fimV* mutant was not able to perform the twitching motility (Fig. 4), as expected (Semmler *et al.,* 2000). Initiation of biofilm formation was also analyzed after five hours of adhesion of cells on a silicon slide at the air-liquid interface. The cultures were adjusted to an OD_600_ = 0.05 in M63 minimum medium supplemented with glucose and casamino acids. The silicon slide was placed upright in a culture tube and cells at the air-liquid interface were analyzed by field emission scanning electron microscope (FESEM), after different time points (Fig. 4 D). PA14 early biofilm presented an irregular architecture due to the motility of the initial adhering cells, but only round microcolonies that did not expand on the surface were observed for the *ΔdgcP* mutant (Fig. 4 D). These results show that DgcP is important to early stages of biofilm formation and twitching motility.

**Figure 4.**
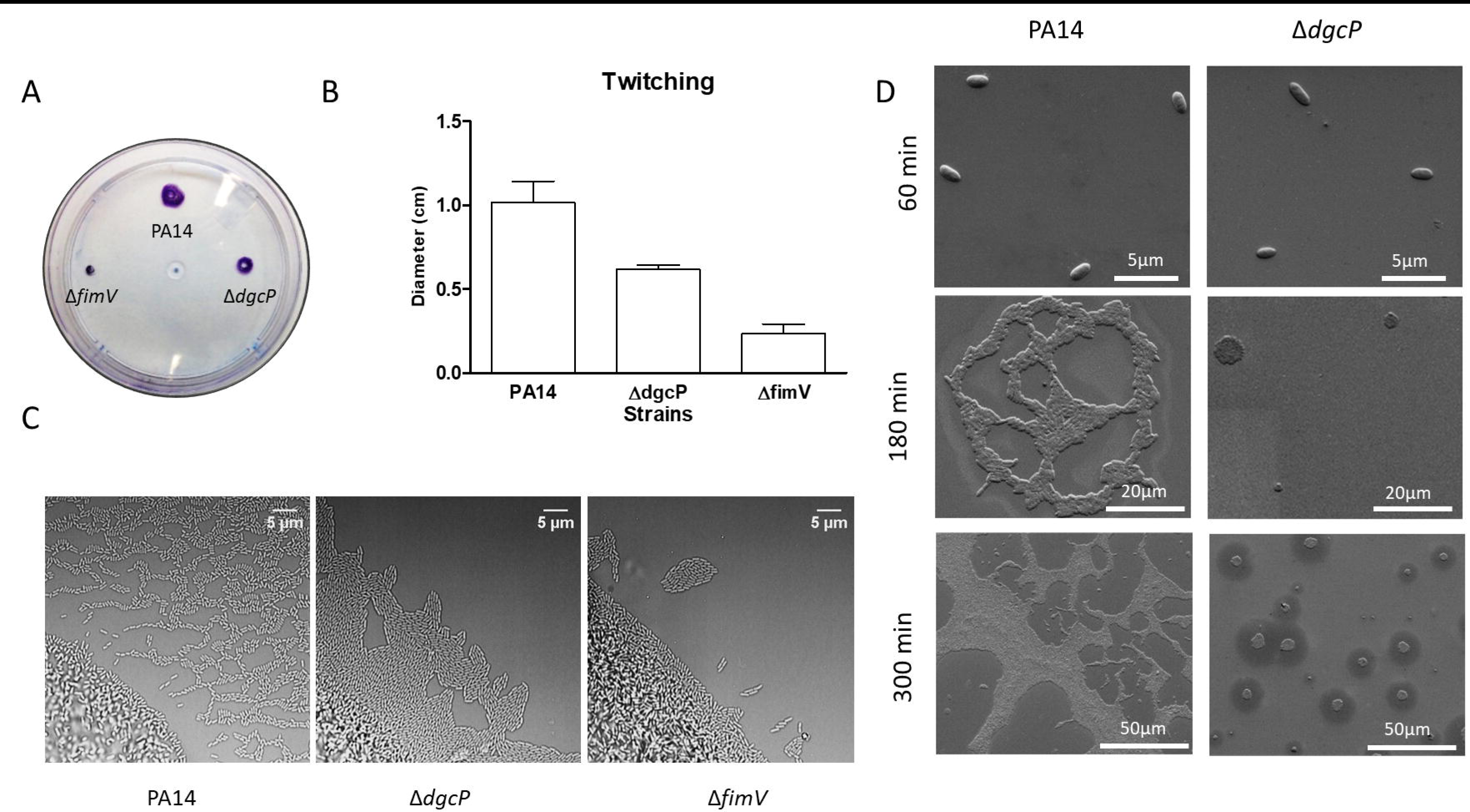
ΔdgcP mutant presents defects related to surface behaviors. Cells were stabbed into the bottom of an agar plate by using a toothpick and incubated upright at 37°C overnight, followed by 48 h of incubation at room temperature. The medium was discarded and adherent cells stained with crystal violet (A). Diameter of the twitching colonies was measured in triplicates. Data are the means ± SD (B). Light microscopy images of PA14 and *ΔdgcP* twitching colonies. Interstitial biofilms formed at the interface between a microscope slide coated in solidified nutrient media (Gelzan Pad) after four hours of colony expansion. The *ΔfimV* mutant was used as a negative control of twitching (C). A silicon slide was placed upright in a culture tube and after the different time points cells at the air-liquid interface were washed, fixed and the spread of cells during the initial stages of biofilm formation was observed by FESEM (D).

### DgcP activity is FimV dependent

Here we observed that the FimV protein localizes the diguanylate cyclase DgcP at the cell pole (Fig. 2). Thus, we asked whether DgcP activity could be regulated by FimV. To answer this question, we overexpressed the DgcP-msfGFP in the *ΔfimV* mutant and analyzed the phenotypes related to c-diGMP. Overexpression of the DGCs DgcP and WspR fused to msfGFP in PA14 increases biofilm formation and decreases swimming motility, indicating that these fusions are functional. Both DGCs also complement the *ΔdgcP* mutation, but DcgP-msfGFP is not able to increase biofilm formation or decrease swimming motility in the *ΔfimV* background, suggesting that it needs FimV for full activity. This is not observed for WspR-msfGFP, which has the same effect with or without FimV in the cells. Overexpression of the C-terminal portion of DgcP (pDgcP-Cterm) that does not localize to the cell poles has no effect on biofilm formation and swimming motility (Fig. 5 A and B) in all strains tested, even though there is an increase in overall c-di-GMP levels (Fig. 5 C), suggesting that localization of the diguanylate cyclase activity is important for those phenotypes. Moreover, overexpression of a mutated DgcP in the diguanylate cyclase motif (GGEEF to GGAAF) decreases biofilm formation in the wild type PA14, but has no effect on both *ΔdgcP* and *ΔfimV* backgrounds (Fig. 5 A and B). DGCs are dimeric proteins therefore the GGAAF mutation may act as a negative dominant on the wild type DgcP.

CdrA is a extracellular protein considered as a scaffold for the biofilm extracellular matrix and transcription of *cdrA* is c-di-GMP-dependent, via FleQ (Borlee *et al.,* 2010) and it is widely used as a reporter of c-di-GMP levels (Rybtke *et al.,* 2012). Overexpression of DgcP-msfGFP leads to ~10-fold increased *cdrA* mRNA levels in PA14 and *ΔdgcP,* but only fourfold in the *ΔfimV* strain (Fig. 5 D). Quantitation of c-di-GMP agrees with the *cdrA* expression levels (Fig. 5 C). Exopolysaccharide (EPS) is also an indication of c-di-GMP levels in several bacteria (Chen *et al.,* 2014; Reichhardt *et al.,* 2015). DgcP and FimV were overexpressed alone or in combination in *Escherichia coli* cells and the production of EPS was assessed in Congo red plates. FimV overexpression does not result in EPS production, but colonies overexpressing DgcP have a pale tint. When both proteins are overexpressed together, EPS production increases, resulting in pink colonies (Fig. 5 E). Altogether, our results corroborate the hypothesis that the polar localization of DgcP by FimV also regulates its activity and that DgcP may contribute to a local c-di-GMP pool.

## Discussion

Recently, Aragon and collaborators showed that DgcP is a well conserved DGC protein in Pseudomonads related to plant and human infections (Aragon *et al.,* 2015). Previously, we showed that overexpression of this protein alters biofilm formation, swimming and swarming motilities as well as imipenem fitness, due to reduced levels of OprD (Nicastro *et al.,* 2014). However, we could not conclude that those phenotypes were specifically related to the physiological role of DgcP, because it was assumed that the overexpression of a DGC increases the global c-di-GMP levels. Herewith, we used protein-protein interactions, characterization of a deletion mutant and protein localization to look for the specific function of DgcP.

*P. aeruginosa* possesses polar T4P which are used for twitching motility and adhesion (Burrows, 2012), essential traits for mature biofilm architecture. Assembly of T4P at normal conditions requires FimV (Wehbi *et al.,* 2011), which shares similar domain organization with *Vibrio cholerae* HubP. These proteins, despite low overall sequence similarity, present a conserved N-terminal periplasmic domain required for polar targeting, and a highly variable C-terminal acidic cytoplasmic region, implicated in protein-protein interactions. HubP is required for polar localization of the chromosomal segregation and chemotactic machineries (Yamaichi *et al.,* 2012; Rossmann *et al.,* 2015). We found that DgcP is present only at the cell poles and that this pattern is dependent on FimV. Thus, we hypothesized that DgcP localization is important for the formation of a localized c-di-GMP pool at the cell poles that would assist the assembly and/or function of the T4P apparatus or other pole-localized organelles (Fig. 6). We suggest that DgcP could be one of the sources of c-di-GMP required for pilus biogenesis in *P. aeruginosa* and probably in other gamma-proteobacteria that carry DgcP and FimV homologs.

**Figure 6.**
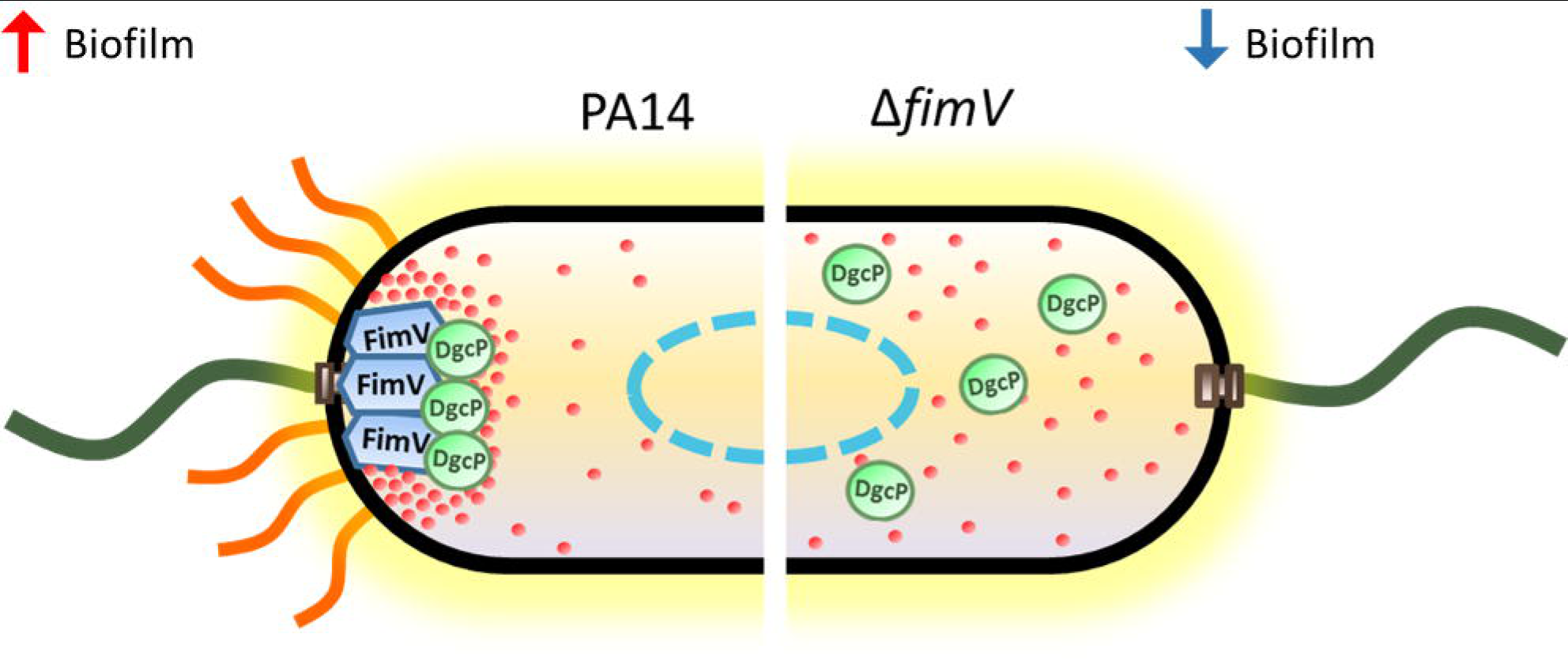
Model of FimV-dependent localization and activity of DgcP. In the PA14 wild type strain, DgcP is located at the pole due to interaction with FimV and may contribute to a local c-di-GMP pool (left). In a *ΔfimV* background, DgcP is scattered in the cytoplasm and may have a small contribution to the global c-di-GMP pool (right). FimV, blue boxes; DgcP, green elipses; c-di-GMP, red dots; nucleoid, blue dashed line; T4P, orange wavy lines and flagella, dark green lines.

Assembly of *P. aeruginosa* T4P at normal conditions requires FimX, a polarly localized c-di-GMP binding protein (Jain *et al.,* 2012) that has degenerate DGC and PDE domains and seems to be enzymatically inactive (Navarro *et al.,* 2009). It is possible that binding of c-di-GMP to the EAL domain of FimX implicates it as an effector protein rather than a PDE, and a FimX mutant that does not bind c-di-GMP fails to activate PilB and twitching motility (Kazmierczak *et al.,* 2006). The FimX homolog in *Xanthomonas citri* interacts with a PilZ protein required for surface localization and assembly of pilin, but does not bind c-di-GMP (Guzzo *et al.,* 2013). X. *citri* PilZ subsequently interacts with PilB, an ATPase required for T4P polymerization, in a cascade of protein-protein interactions (Guzzo *et al.,* 2009; Dunger *et al.,* 2014).Remarkably, suppressor mutations in a *P. aeruginosa fimX* mutant that restored T4P biogenesis and partially restored twitching motility also increased c-di-GMP levels. However, the suppressor mutant cells presented peritrichous pili (Jain *et al.,* 2012), suggesting that, in addition to FimX, a more specific source of c-di-GMP would be needed for the correct assembly of the machinery at the cell poles. Similarly, a *P. aeruginosa* PilZ domain protein is also involved in the T4P-based twitching motility and does not bind to c-di-GMP, suggesting a conserved mechanism (Burrows, 2012). Cumulatively, these findings imply that the molecular mechanisms of pilus protrusion and retraction are regulated by local fluctuations of c-di-GMP levels. Other components of polar localized structures, such as flagella and the chemotactic machinery also bind directly or indirectly to c-di-GMP (Düvel *et al.,* 2012; Baker *et al.,* 2016), and DgcP may have a function in those mechanisms as well, but further work would be needed to uncover such roles.

*P. aeruginosa* T4P was demonstrated to be important not only to attach and move, but also to sense mechanical features of the environment. T4P sensing on solid surface increases its extension and retraction frequencies and cAMP production, leading to the upregulation of the cAMP/Vfr-dependent pathway (Persat *et al.,* 2015). Recently, FimV was associated with this process by interaction with FimL, a scaffold protein that connects T4P with the Chp chemosensory system via interaction with PilG and FimV (Inclan *et al.,* 2016; Buensuceso *et al.,* 2016). FimV is also involved in the transcriptional upregulation of *pilY1,* and PilY1 increases the SadC diguanylate cyclase activity upon surface contact (Luo *et al.,* 2015).However, SadC presents localized foci at the poles, in the middle of cells and between these two locations (Zhu *et al.,* 2016), suggesting that it may have a broader role. Thus, we suggest that an outside signal could be sensed by T4P and transduced by FimV as described for SadC, resulting in the direct and localized activation of c-di-GMP production by DgcP. The finding that FimV-DgcP interact at the poles is an important step towards the understanding of how c-di-GMP localized pools are formed, controlling the spatial activity of target proteins.

## Experimental procedures

### Bacterial strains, plasmids and growth conditions

The bacterial strains and plasmids used in the study are described in the Supplementary Table S2. For routine cell cultures, bacteria were grown aerobically in Luria-Bertani (LB) broth or LB agar at 37 or 30°C. Ampicillin (100 μg/ ml), kanamycin (50 μg/ml) or gentamicin (10 μg/ml) were added to maintain the plasmids in *E. coli.* Carbenicillin (300 μg/ ml), kanamycin (250 μg/ml) or gentamicin (50 μg/ml) were added to maintain the plasmids in *P. aeruginosa.* For the pJN105 related constructs (Table S1), arabinose was added to cultures at 0.2 % final concentration. Both M8 (Kohler *et al.,* 2000) and M63 (Pardee *et al.,* 1959) minimal salt media were supplemented with 1mM MgSO_4_, 0.2 % glucose and 0.5 % casamino acids (CAA). To visualize bacterial two-hybrid interactions on solid medium, MacConkey indicator medium (Difco) supplemented with 1 % maltose and 100 mM IPTG (Isopropyl β-D-1-thiogalactopyranoside), herein designated MacConkey medium, was used.

### General molecular techniques

DNA fragments were obtained by PCR using Q5 DNA polymerase (NEB). Oligonucleotide primers were purchased from Life Technologies and the sequences are listed in Table S1. PCR products of the expected sizes were purified from gels using GeneJET TM Gel Extraction Kit (Thermo Scientific), cloned using the SLIC method (Jeong *et al.,* 2012) and transformed into *E. coli* DH5α (Table S1). Plasmid purification was performed with GeneJET Plasmid Miniprep kit (Thermo Scientific). Sequencing was carried out using the Big Dye terminator cycle sequencing kit (Applied Biosystems) using the facility of the Departamento de Bioquímica, IQ-USP (SP, Brazil).

To construct unmarked in-frame deletions, the upstream and downstream regions of the target gene were amplified and cloned into the pEX18Ap *(fimV)* or pKNG *(dgcP).* The resulting constructs were used to delete target genes on wild type PA14 genome by homologous recombination. To construct the pDgcP plasmid the *dgcP* coding region was cloned in frame with a synthetic *msfGFP* gene into the pJN105 plasmid. The *msfGFP* codes for a N-terminal 40 amino acids spacer and a C-terminal monomeric super fold GFP. All vectors and constructs are described in more detail in Table S1.

### Biofilm assays

Three different biofilm assays were performed. The microtiter dish biofilm formation assay was performed as described (O’Toole, 2011). The biofilms observed by confocal laser scanning microscopy (CLSM) were grown in 8-well chamber slides and stained with DAPI as described (Jurcisek *et al.,* 2011) and imaged using a Zeiss-LSM 510-Meta. For the early stages of biofilm formation on silicon wafers, cultures were adjusted to an OD_600_ = 0.05 in M63 medium and transferred to a 24-well plate where silicon slides (21 cm^2^) were placed upright in each well. Before using the silicon substrates, they were previously cleaned by ultra-sonication for a period of 15 min each in acetone, isopropanol and distilled water, respectively. Slides were dried under N_2_ flow and subsequently treated with O_2_ plasma at 100 mTorr for 15 min (720 V DC, 25 mA DC, 18 W; Harrick Plasma Cleaner, PDC-32G). After 60, 180 or 300 minutes the slides were rinsed three times with water, fixed with 4% paraformaldehyde for 1h and analysed by field-emission scanning electron microscopy (FESEM; model F50, FEI Inspect) operated at 2 keV. Prior to examination, samples were coated with sputtered gold to prevent electrical charging.

### High-throughput two-hybrid assays

PAO1 two-hybrid library (Houot *et al.,* 2012) was tested against the pKT25_DgcP bait. Basically, 25-50 ng of pUT18 library was transformed into BTH101 cells carrying the pKT25_DgcP vector and plated on MacConkey medium for 48-96 h at 30 °C. Red colonies were picked up and restreaked on MacConkey plates. The positive colonies were cultivated in liquid medium, and plasmids were isolated and further analyzed. The candidate preys were retested individually for interaction with the bait by retransforming pUT18 derivative prey and pKT25 bait plasmids into BTH101 cells and also the pUT18 derivative preys and the pKT25 empty vector. The interaction was evaluated by the color of spotted co-transformants on MacConkey plates and β-galactosidase assays. Cells were grown on MacConkey plates for 96 hours and they were scrapped and suspended in 1mL of PBS. 100 μL were used in the classical β-galactosidase assay (Miller, 1972).

### Twitching assay

Macroscopic twitching assay was performed as described (Dae-Gon Ha *et al.,* 2014) with minor modifications. Briefly, a colony was picked with a toothpick and stabbed at the bottom of a plate containing M8 medium supplemented with 1 mM MgSO_4_, 0.2 % glucose, 0.5 % casamino acids and 2.0 % agar. The plates were incubated upright at 37°C overnight, followed by 48 h of incubation at room temperature (~25°C). Next, the agar was removed, and the bacteria were stained with 0.1% crystal violet. Microscopic twitching assay was performed as described (Turnbull and Whitchurch, 2014).

### Swimming assay

Swimming assays were performed by inoculating 5Lf.il of a stationary phase-grown liquid cultures in M8 with 0.3% agar that were incubated for 16Lh at 30°C in a plastic bag to maintain the humidity constant (Dae Gon Ha *et al.,* 2014).

### Congo red assay

5Lf.il of stationary phase-grown cultures were inoculated at 1% agar plates of tryptone broth (10 g/L) containing Congo red (40 μg/mL) and Coomassie brilliant blue (20 μg/mL). The plates were incubated for 16 h at 30°C and then for 96 hours at room temperature.

### qRT-PCR

For qRT-PCRs, total RNA was extracted with Trizol (Invitrogen), treated with DNase I (Thermo Scientific, Waltham, MA, USA) and used for cDNA synthesis with Improm II (Promega) or Superscript III (Invitrogen) and hexamer random primers (Thermo Scientific). cDNA was then amplified with specific primers using Maxima SYBRGreen/ROX qPCR Master Mix (Thermo Scientific) and the 7300 Real Time PCR System (Applied Biosystems). *nadB* was used as internal control for normalization of total RNA levels (Lequette *et al.,* 2006). The relative efficiency of each primer pair was tested and compared with that of *nadB* and the threshold cycle data analysis (2'^AACt^) was used (Livak and Schmittgen, 2001). All reactions were performed in triplicates, the assays were repeated at least twice using independent cultures and the results of one representative experiment are shown, with average values of technical triplicates and error bars representing standard deviation of ΔΔCt.

### Fluorescence and light microscopy

To verify the localization of DgcP_msfGFP fusions, fluorescence microscopy was performed using a Nikon Eclipse TiE microscope equipped with a 25-mm SmartShutter and an Andor EMCCD i-Xon camera. For fluorescence microscopy and bright field microscopy, a Plan APO VC Nikon 100X objective (NA = 1.4) and a Plan Fluor Nikon 40X objective (NA = 1.3) were used. For membrane staining, cells were treated with 50 μg/mL FM4-64 (Invitrogen). For phase contrast microscopy, a Plan APO A OFN25 Nikon 100X objective (NA = 1.45) was used. All microscopy assays were performed with immobilized cells on 25 % LB pads with 1.5% agarose. Image analyses were performed using the ImageJ (Schneider *et al.,* 2012) and MicrobeJ (Ducret *et al.,* 2016) softwares.

### c-di-GMP extraction and quantification

c-di-GMP was extracted as described by (Irie and Parsek, 2014) with minor modifications. 50 mL of cultures were grown in M8 medium at 37°C and 200 rpm until reach OD.600 = 1. Cells were collected by centrifugation resuspended in 500 μL of M8 medium with 0.6 M perchloric acid. The tubes were incubated on ice for 30 minutes and then centrifuged at 20000 *g* for 10 minutes. The pellets were used for protein quantitation and the supernatants were neutralized with 1/5 volume 2.5 M KHCO3. The nucleotide extracts were centrifuged again and the supernatants were stored at-80 °C. High-performance liquid chromatography (LC) was performed using the 1200 Infinity LC System (Agilent) that consists of a degasser, a quaternary pump, a thermostated autosampler (4°C) and a temperature (30°C)-controlled column compartment. This system was coupled to an 3200 Qtrap LC-MS/MS system equipped with an Electrospray Ionization source (ESI) (AB Sciex, USA). Analyst 1.4.2 software (AB Sciex, USA) was used to operate the equipment and calculate c-di-GMP concentrations.

Samples (injection of 10 μL) were separated by a Phenomenex Synergi Hydro-RP column (150L*L2 mm, 4 μm) using 0.1% formic acid in 15 mM ammonium acetate as mobile phase A and MeOH as mobile phase B at a flow rate of 0.3 mL min^-1^. The gradient program was 0 min 2 % B, 0.5 min 2 % B, 4.5 min 30 % B, 6.0 min 80 % B, 7.0 min 80% B, 7.01 min 2% B and 14 min 2 % B. For quantification of c-di-GMP, the tandem mass spectrometry method multiple reaction monitoring (MRM) was used in negative mode. The following parameters were set: nebulizer, heated auxiliary and curtain gases (nitrogen) at 20, 30, 10, respectively; Turbo IonSpray voltage and temperature at-3,800 V and 250 °C, respectively; MRM transition (in m/z) 689.1 → 344.2 with a dwell time of 200 ms per transition; collision energy (CE) at-45 eV; and declustering potential at-53 V. An external standard curve was prepared for c-di-GMP in the MRM mode. The stock solution was diluted and the c-di-GMP peak area plotted against the nominal concentrations (16 to 2,000 ng mL-1).

### Multiple Sequence Alignment and secondary structure prediction

Reciprocal best-hit search was performed as described before (Kohler et al., 2015). Briefly, the Kyoto Encyclopedia of Genes and Genomes (KEGG) database was initially used to search open-reading frames showing highest identity against DgcP (GenBank: ABJ14873.1). This search resulted in a list that was used to reciprocal search its highest identity orthologue in *P.aeruginosa* UCBPPA14 genome. In both searches the threshold for Smith-Waterman score was 100, and the resulting best-best hit was filtered using a minimum 20% identity comparing the identified homologue against the DgcP. A list of all organisms that displays a DgcP homologue are listed in Supplemental Table S1. Redundant sequences were filtered using CD-Hit (Li and Godzik, 2006) with a maximum identity of 90% (Table S1). A multiple sequence alignment was performed using MUSCLE software (Edgar, 2004). The multiple sequence alignment was then visualized in the jalview (Waterhouse et al., 2009) and secondary structure analysis predicted with JPred (Drozdetskiy et al., 2015).

## Acknowledgments

We would like to thank A. Bisson-Filho for kindly providing the msGFP synthetic gene, D. Schechtman and M. Navarro for carefully reading the manuscript. We are also in debt with F. Gueiros-Filho, L. Zambotti-Villela and M.A. Cotta for assistance with fluorescence microscopy, mass spectrometry and electron microscopy, repectively. We acknowledge the National Nanotechnology Laboratory (LNNano, CNPEM) for granting access to the electron microscopy facilities.

Conceived and designed the experiments: GGN, CB and RLB. Performed the FESEM experiments: JHM. Performed two-hybrid screening: GGN and CB. Performed fluorescence and light microscopy: GGN and AAP. General molecular procedures: GGN, GHK, ALB and TOP. HPLC-MS/MS: ES. Contributed reagents/materials/analysis tools: RLB, CB, PC. Wrote the paper: GGN, RLB.

G.G.N. is supported by São Paulo Research Foundation (FAPESP) grant numbers: 2013/02375-1 and 2014/02381-4.

R.L.B. is partially supported by National Council for Scientific and Technological Development (CNPq 307218/2014-7) a São Paulo Research Foundation (FAPESP 2014/05082-8).grant allowed the work in RLB laboratory.

C.B. is supported by the ANR grants REGALAD ANR-14-CE09-0005-02.

The authors declare that there is no conflict of interest.

---

**Figure S1. Multiple Sequence Alignment (MSA) and secondary structure prediction of DgcP homologues.** The MSA was generated using Muscle and edited in Jalview. The amino acids residues are colored using the Clustal X colour scheme. Secondary structure prediction was generated using JPred and is displayed as alpha-helixes (red cylinders) and beta-sheets (green arrows) on the top of the alignment. Horizontal black bars represent the two putative N-terminal domains and the blue bar shows the GGDEF C-terminal domain. The orange vertical bar on the left point to *Pseudomonas* spp. and the purple bar groups other bacteria, as listed in **Table S1**.

